# Linking impulsivity to activity levels in pre-supplementary motor area during sequential gambling

**DOI:** 10.1101/2022.06.28.497876

**Authors:** Allan Lohse, Annemette Løkkegaard, Hartwig R. Siebner, David Meder

## Abstract

Impulsivity refers to the tendency to act prematurely or without forethought, and excessive impulsivity is a key problem in many neuropsychiatric disorders. Since the pre-supplementary motor area (preSMA) has been implicated in inhibitory control, this region may also contribute to impulsivity. Here, we examined whether functional recruitment of preSMA may contribute to risky choice behavior (state impulsivity) during sequential gambling and its relation to self-reported trait impulsivity. To this end, we performed task-based functional MRI (fMRI) after low-frequency (1 Hz) repetitive transcranial magnetic stimulation (rTMS) of the preSMA. We expected low-frequency rTMS to modulate task-related engagement of the preSMA and hereby, tune the tendency to make risky choices. 24 healthy volunteers (12 females, 19-52 years) received real or sham rTMS on separate days in counterbalanced order. Thereafter, participants performed a sequential gambling task with concurrently increasing stakes and risk during whole-brain fMRI. In the sham-rTMS session, self-reported trait impulsivity scaled positively with state impulsivity (riskier choice behavior) during gambling. The higher the trait*-*impulsivity, the lower was the task-related increase in preSMA activity with increasingly risky choices. Following real-rTMS, low-impulsivity participants increased their preference for risky choices, while the opposite was true for high-impulsivity participants resulting in an overall decoupling of trait impulsivity and state impulsivity during gambling. This rTMS-induced behavioral shift was mirrored in the rTMS-induced change in preSMA activation. These results provide converging evidence for a causal link between the level of task-related preSMA activity and the propensity for impulsive risk-taking behavior in the context of sequential gambling.

**Significance statement:** Impulsivity is a personal trait characterized by a tendency to act prematurely or without forethought, and excessive impulsivity is a key problem in many neuropsychiatric disorders. The pre-supplementary motor area (preSMA) has been implicated in inhibitory control. Here, we provide evidence that this region contributes to the implementation of general impulsive tendencies (trait impulsivity) into actual behavior (state impulsivity). When healthy volunteers performed a sequential gambling task, their choice behavior (i.e., state impulsivity) correlated positively with their impulsivity score (i.e, trait impulsivity). Additionally, participants with lower trait impulsivity showed a stronger increase in task-related activity of the preSMA with increasing risk. Both of these relationships were uncoupled after perturbing the preSMA with repetitive transcranial stimulation (rTMS).

## Introduction

Impulsivity is a personal trait that describes one’s tendency to act prematurely or without forethought (Dalley and Robbins, 2017). Excessive impulsivity is intimately linked to a breakdown of inhibitory processes (Bari and Robbins, 2013) and constitutes a prominent clinical feature in many neuropsychiatric conditions (Kulacaoglu et al., 2017; Kozak et al., 2019). Trait impulsivity has multiple facets which are commonly assessed with self-reports reflecting attentional, motor, and planning aspects of impulsivity (Stanford et al., 2009). Specific facets of impulsivity can also be experimentally quantified with tasks probing the inability to suppress premature response tendencies (*waiting impulsivity*) (Voon et al., 2014), the failure to cancel already initiated responses (*stopping impulsivity*) (Logan et al., 1984) or the preference for larger, less certain rewards over smaller, more certain rewards (*risky impulsivity*) (Robbins and Dalley, 2017). While these tasks yield a momentary read-out of the impulsive state of the individual at the time of examination (state impulsivity), behavioral measures of impulsivity often correlate only weakly with self-report trait measures of impulsivity and every-day risk-taking behavior (Schonberg et al., 2011; Cyders and Coskunpinar, 2012). Thus, there is little knowledge about whether and how trait impulsivity might mediate state impulsivity assessed in the laboratory.

Task-related functional MRI (fMRI) has been used extensively to identify brain networks that are implicated in impulsive behaviors. These studies have consistently reported altered cortical activity in the pre-supplementary motor area (preSMA) and right inferior frontal cortex (rIFG), in healthy individuals as well as patients suffering from psychiatric disorders with impaired impulse control (Taylor et al., 2007; Forstmann et al., 2008; Tabibnia et al., 2011; Ersche et al., 2012; Whelan et al., 2012). Together with electrophysiological studies, the fMRI studies corroborate the notion that the rIFG and preSMA, together with the subthalamic nucleus (STN), form a network supporting fast and non-selective suppression of inappropriate action tendencies, often referred to as the braking network (Aron et al., 2007; Wessel et al., 2019). Given the correlative nature of functional brain mapping, it is unclear whether and how the neural structures that have been associated with impulsivity are causally involved in the expression of impulsive actions.

Using task-related fMRI, we showed that the preSMA, IFG, and STN increase their activity during a sequential gambling task in which individuals either continue to gamble or stop in the context of increasing stake (Meder, Haagensen et al., 2016). These areas showed a gradual increase in activity with increasing stake which was paralleled by a gradual increase in reaction time. Based on these findings, we argued that the braking network does not only generate an acute stop signal to pause ongoing actions, but also implements a gradually increasing braking signal that prevents impulsive choices during sequential gambling. However, our fMRI findings are only correlational in nature, and we did not test for a relationship between the variability in gambling behavior (state impulsivity) and the subject’s self-reported trait impulsivity.

Following-up on our fMRI study (Meder, Haagensen et al., 2016), we used a perturb-and-map approach to probe the causal contribution of the preSMA in the control of choice impulsivity during sequential gambling (Bergmann et al., 2016). We first targeted the preSMA with slow-frequency (1 Hz) repetitive transcranial magnetic stimulation (rTMS) using robot-assisted neuronavigation. We then performed fMRI to probe the impact of rTMS on task-related activity in preSMA and risky choice behavior during gambling (state impulsivity), using the sequential gambling task introduced by Meder, Haagensen et al. (2016). We also assessed trait impulsivity with the Barratt impulsiveness scale (BIS-11) scale.

We hypothesized that trait impulsivity would predict inter-individual variations in the tuning of task-related activation of preSMA to gradual increases in stake as well as inter-individual differences in choice behavior during the gambling task. We further predicted that the functional perturbation evoked by 1Hz rTMS over the preSMA would modulate choice impulsivity expressed during the sequential task and that the modulatory effect of rTMS would depend on the participants’ trait impulsivity.

## Materials and methods

### Participants

We included 24 healthy volunteer adults (mean age 28.2 years; SD 9.7 years, range: 19-52 years; 12 females) with no history of mental or neurological illness. One participant was excluded from the analysis since this participant later revealed not having fully understood the task instructions for the first session, which was also clear from the change in the task performance between the two sessions. One participant was only included in the behavioral analysis, since the data of one fMRI session was not saved due to an error.

The study was approved by the research ethics committee of the capital region of Denmark (H-15017878) and all participants gave written informed consent prior to participating in the study.

### Experimental design

The sequential gambling task was a computerized, open-ended, one-player version of the dice game “pig” first described by the American magician John Scarne (Scarne, 1945; Meder, Haagensen et al., 2016, Fig. 1). Each round began with a die being rolled automatically (the “rolling” was visualized by a rapid succession of one of the six sides of the die in random order (presentation time 150 ms)) for a jittered period of time (roll time 1.5-3.5 s). After observing the outcome of the throw, the participant had two seconds to decide to roll again and accumulate more points or to hold and bank the accumulated points. The participant could roll the die as many times as he or she wanted. The round ended when the participant holds and thereby banked the sum of the rolls during that round or if he or she rolled a 1 in which case the accumulated sum was lost and a new round began. The outcome of the round was shown to the participant for 2.5 seconds. The pay-out of the game was the average sum accumulated in all rounds, including loss rounds, multiplied by 10 Danish kroner (DKK; 1 DKK ≈ 0.15 USD). Every six minutes, participants switched between two types of task blocks which differed in terms of motor engagement. In “act-to-continue” blocks, participants had to actively press a button to continue with gambling, and to refrain from a button press to end the gambling round in order to receive the accumulated sum. In “act-to-stop” blocks, sequential gambling did not require any action as the die was rolled automatically until participants actively ended the round by pressing a button. Between blocks, participants saw an instruction screen for 60 seconds. In this paper, the different action contexts of gambling were not considered in the analysis. This aspect of the task will be addressed in a separate paper.

**Figure 1.**
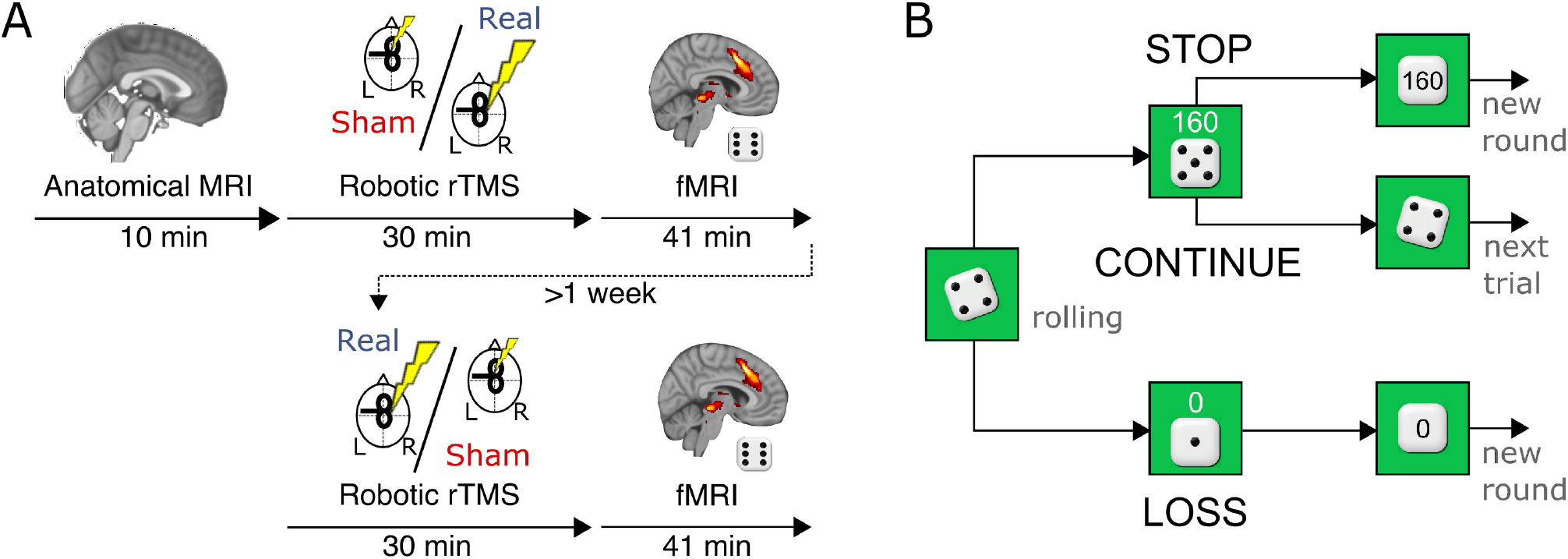
Task and experimental setup. (A) A total of 24 healthy volunteers participated in a two-day study. On two days separated by at least one week, participants underwent 30 minutes of either inhibitory (1 Hz, 100% rMT) repetitive transcranial magnetic stimulation (rTMS) of the right presupplementary motor area (preSMA) or a sham stimulation (right preSMA, 1 Hz, 30 % rMT). The order of the stimulations was randomized and counterbalanced. Immediately following rTMS, participants performed a gambling task during fMRI recording to assess neural activity. (B) In the gambling task, participants accumulated points as they repeatedly rolled a die until they decided to bank the winnings and start a new round, or until rolling a “1” resulting in the loss of all points in this round. Total winnings were the average of all rounds, including loss rounds. *rMT*, resting motor threshold.

The known, true odds for not losing every time a die is rolled are 5:1. The average winning amount is 40 DKK 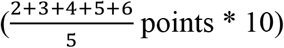. Thus, if the accumulated sum is less than 200 DKK (resulting in odds 200:40 ∼ 5:1), the odds for winning are in favor of continuing rolling (Knizia, 2001). Thus, for a gain-maximizing strategy, participants should always stop the round once having accumulated 200 or more points.

### Repetitive Transcranial Magnetic Stimulation

Participants underwent real-rTMS (100 % resting motor threshold; rMT) and sham-rTMS (30 %rMT) rTMS of preSMA in two separate sessions at least one week apart. The session order was counterbalanced across participants. For both sessions, over the course of 30 minutes 1800 biphasic pulses were applied over preSMA with a frequency of 1 Hz using a robot arm-controlled MCF-B65 figure-8 coil (Ø: 2 × 75 mm; Medtronic). The coil direction was positioned in a lateral-to-medial orientation such that the strongest intracranial current went in a left-right direction, thus preferentially targeting the right hemisphere.

The individual resting motor threshold was determined in all sessions using the freeware TMS Motor Threshold Assessment Tool version 2.0 (MTAT 2.0, http://www.clinicalresearcher.org/software.htm) where the threshold is estimated using a maximum-likelihood strategy (“threshold hunting”). The rMT was assessed in the right hemisphere and the motor evoked potential (MEP) of the left first dorsal interosseous was used as read-out using the same coil type as the one used during the experiment. MEPs had to exceed 50 μV to be considered present.

### Robot-assisted neuronavigation

Right preSMA was localized using neuronavigation (LOCALITE, Sankt Augustin, Germany) based on individual T1-weighted MRI scans acquired prior to the first TMS session. After normalizing the participants’ anatomical scans into standard MNI space the stimulation target was set to [6, 16, 52] based on the peak activation from the previous study done at our center using the same task (Meder, Haagensen et al., 2016). For the duration of the stimulation the coil position was automatically maintained by a TMS robot (Axilum Robotics, Strasbourg, France). After the stimulation the participants were transferred to the scanner in an MR compatible wheelchair.

### Structural and Functional MRI

Scans were acquired using a 3T Verio scanner with a 32-channel head coil (Siemens, Erlangen, Germany). To relate the blood oxygen level dependent (BOLD) signal acquired with fMRI to anatomical brain structures and for rTMS neuronavigation a structural T1-weighted image was acquired (MPRAGE; repetition time [TR] 1900 ms, echo time [TE] 2.32 ms, flip angle 9°in-plane resolution .89×.89 mm; slice thickness .9 mm; field of view [FOV] 256 mm). To assess task-related neural activity, regional blood-oxygen-level dependent (BOLD) signal changes were measured using a T2*-weighted EPI sequence (TR 1650 ms, TE 26 ms, flip angle 74°). 1459 volumes with 32 slices in ascending order were acquired per session (in-plane resolution 3×3 mm; FOV 192 mm). Axial slices were arranged parallel to the bicommissural line. Respiration and pulse were obtained with a pneumatic thoracic belt and pulse oximeter, respectively.

### Preprocessing of functional MRI data

To allow for the T1 equilibrium effect the first three volumes of each run were discarded. The EPI volumes of each session were slice time corrected, realigned to the first volume in the time series and unwarped with FSL topup tool. The unwarped images from the two sessions of each participant were then realigned to the mean image, segmented, and co-registered to the segmented T1-weighted image and normalized to an MNI space template using affine warping and a DCT basis (Ashburner and Friston, 1999) and smoothed with a 8 mm full-width half-maximum (FWHM) gaussian kernel. Volumes acquired during the pause between blocks were excluded (5 × 36 volumes per session).

### Analysis of functional MRI data

The main aim of this study was to interrogate the causal involvement of the increase in preSMA activity during sequential gambling in the mediation of risky decision-making. Our main regressor of interest was thus the parametric modulation of “continue” events with the accumulated sum during that decision. The general linear model included the three main events of the paradigm (“continue”, “stop” and “loss”) and their parametric modulation with the accumulated sum in the trial in both block types (*act to continue* and *act to stop* blocks). Additional regressors of no-interest were onsets of the dice roll, onsets of rounds where the first outcome was a 1, and onsets of the feedback screen for both loss and win rounds. Heart and pulse rate were added to the design matrix as nuisance regressors (Lund et al., 2006). Region of interest (ROI) analyses were performed using a preSMA mask defined as a sphere (r = 5 mm) centered at the stimulation target (MNI coordinate *x, y, z* = [6,16,52]), putamen using the anatomical masks from the WFU pick Atlas (Maldjian et al., 2003) and subthalamic nucleus using a probabilistic mask based on 7T MRI (Keuken et al., 2014). Preprocessing and analysis of the images were performed in SPM12 (revision number 6906, Wellcome Department of Imaging Neuroscience, Institute of Neurology, University College London).

### Statistical analysis

Statistical analyses of behavior were performed in R (R Foundation for Statistical Computing, Vienna, Austria). We used a linear mixed model implemented in the nlme package (Pinheiro et al., 2018) with regressors motor context (act to continue, act to stop), rTMS condition (real-rTMS, sham-rTMS) and session number (session 1, session 2) for analysis of mean stopping amounts. Response times were log-transformed to meet assumption of normal distribution. The mean +/-SD is reported. For correlational analyses Pearson’s *r* is reported.

## Results

### Effects of real-rTMS on choice behavior

Participants completed a mean total of 528.8 *continue* trials, 135.2 *stop* trials and 121.6 *loss* trials. Real-rTMS did not change the mean stopping amount significantly compared to sham-rTMS (mean sham-rTMS 160.40, mean real-rTMS 159.81, paired t-test t(22) = 0.08, p = 0.94). However, in the sham-rTMS session, there was a significant correlation between self-reported impulsivity (BIS-11 score) and mean stopping amount (*r* = 0.43, *p* = 0.040). The higher the individual BIS score, the larger was the mean amount at which participants decided to stop the round and bank their accumulated earnings (mean stop amount). This relationship between self-reported “trait” impulsivity, indexed by the BIS-11 score, and “state” impulsivity, expressed by the participant’s choice behavior during sequential gambling, was no longer present after real-rTMS (*r* = 0.15, *p* =0.48). Accordingly, we found a significant interaction between self-reported impulsivity and rTMS condition for the mean stopping amount (*p* = 0.017; Figure 2B). The relative effect of real-rTMS on mean stopping amount was inversely related to the participants’ trait impulsivity (*r* = −0.49, *p* > 0.001; Figure 2A): The higher the participant’s trait impulsivity, the more did real-rTMS reduce state impulsivity (stopping behavior) relative to sham-rTMS. Conversely, the lower the participant’s trait impulsivity, the more did real-rTMS increase task-related state impulsivity relative to sham-rTMS. Together, these findings indicate that real-rTMS over preSMA led to a decoupling of trait impulsivity and state impulsivity.

**Fig. 2.**
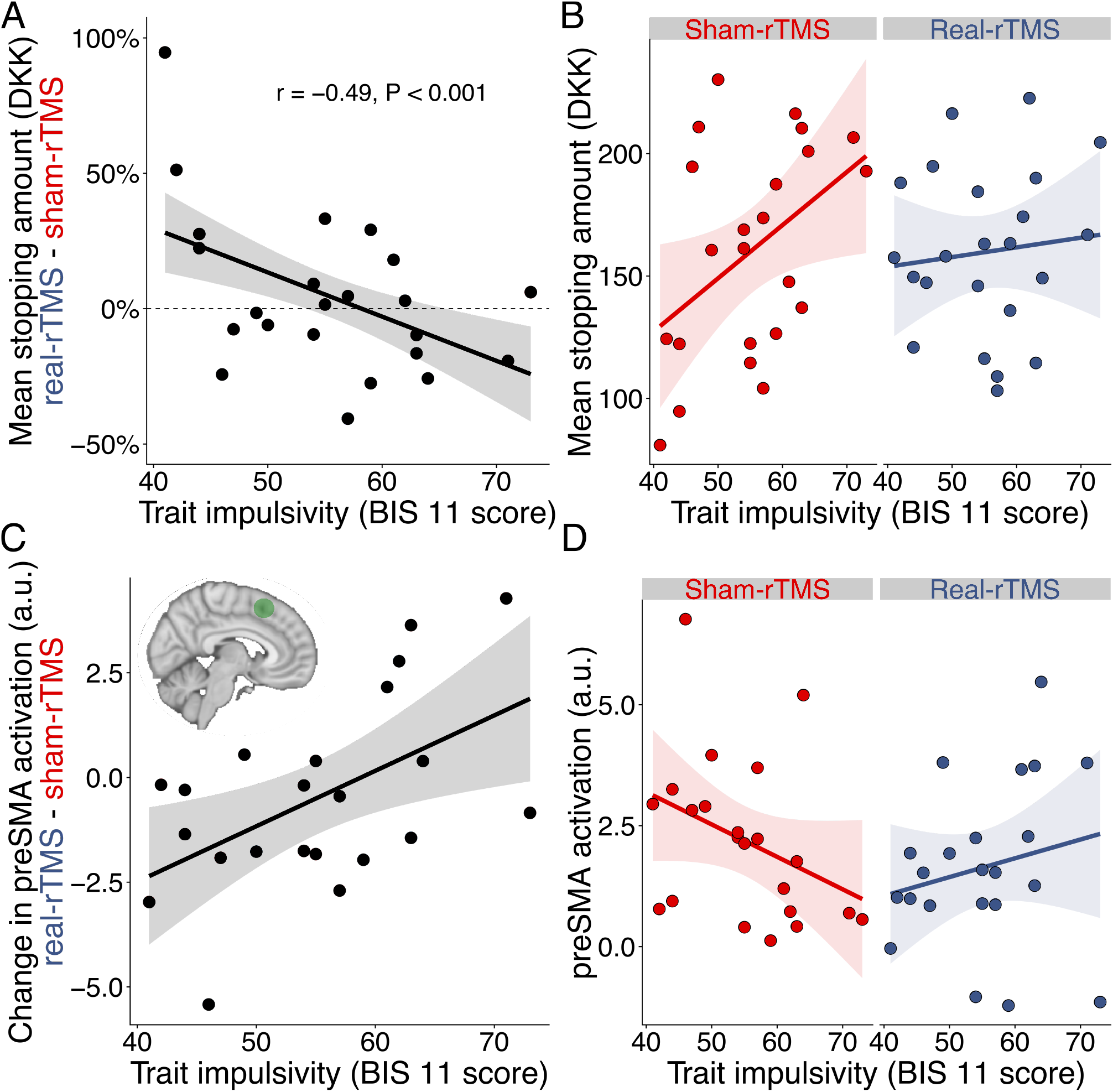
PreSMA causally modulates impulsivity during “continue-to-gamble” decisions. (A) 1 Hz Trait impulsivity predicts rTMS effect on mean stopping amount. Participants who reported low degrees of impulsivity increased their tendency to make risky choices, whereas the opposite was true for those who reported high levels of impulsivity. (B) 1 Hz real-rTMS decouples association between trait impulsivity and task-related risk-taking behavior. In the sham-rTMS condition, impulsivity, measured using a standard inventory (BIS 11), correlated with the mean stopping amount in the dice game. Inhibiting the preSMA with 1 Hz rTMS decorrelated self-reported trait impulsivity and decisional impulsivity during the task. (C) Trait impulsivity predicts the rTMS effect on overall task-related brain activation during continue trials. 1 Hz real-rTMS lowers linear increase in preSMA BOLD signal parametrically modulated by the continue sum and covarying with self-reported trait impulsivity. The region of interest (ROI) was defined as a sphere (r = 5 mm) centered at the stimulation target (MNI coordinate *x, y, z* = [6,16,52]). (D) Overall preSMA activation scaling with accumulated sum following sham-rTMS and real-rTMS. BIS 11, Barratt impulsiveness scale; rTMS, repetitive transcranial magnetic stimulation. *N* = 23 participants

### Response times

Decisions to continue were made more slowly as the accumulated sum increased after both sham-rTMS (mean slope of log-transformed continue RTs: 0.00174, SD: 0.00193; one-sample t(22) = 4.3352, *p* = 0.000266), and real-rTMS of the preSMA (mean slope 0.00172, SD: 0.00152; one-sample t(22) = 5.418, *p* < 0.0001), but there was no difference between the two rTMS conditions (paired t(22) = 0.075978, *p* = 0.9401).

Conversely, decisions to stop gambling and bank the winnings were made more quickly when the stake became higher in both the sham-rTMS (mean slope of log-transformed stop RTs: −0.0025, SD: 0.0026; one-sample t(22) = −4.6303, *p* = 0.0001), and real-rTMS session (mean slope −0.0019, SD: 0.002; one-sample t(22) = −4.5476, *p* = 0.0002), but there was no significant difference between the two rTMS conditions (paired t(22) = 1.09, *p* = 0.29).

### Functional magnetic resonance brain imaging

In the sham-rTMS session, a midline cluster showed a linear increase in “continue-to-gamble” activity with increasingly higher stakes, comprising the preSMA and dorsal ACC (Figure 3; Table 1). The linear scaling of preSMA activity with increasing stakes was inversely related to task-related state impulsivity (Figure 2D): The stronger the “continue-to-gamble” activity in preSMA scaled with the stake, the lower was the participant’s mean stopping amount (Figure 2D). The ventral and dorsal striatum, anterior insula, lateral intraparietal cortex (LIP), inferior parietal lobule, right inferior frontal cortex, subthalamic nucleus and occipital cortex also showed an increase in “continue-to-gamble” activity during a sequential gambling round that scaled with accumulated sum (Figure 3; Table 1), replicating the activity pattern reported in our previous study (citation).

**Table 1.** Significantly activated clusters during sequential gambling.

| Region | z | Peak MNI coordinates: |  |  | z | Peak MNI coordinates: |  |  |
| --- | --- | --- | --- | --- | --- | --- | --- | --- |
|  | score | left hemisphere |  |  | score | right hemisphere |  |  |
|  | peak | x | y | z | peak | x | y | z |
| **SHAM-rTMS SESSION ONLY** |  |  |  |  |  |  |  |  |
| *Task-related activity during “continue” trials showing a linear increase with cumulative gamble sum* |  |  |  |  |  |  |  |  |
| Occipital cortex (C1) | 5.09 | -20 | -94 | -8 |  |  |  |  |
| Anterior insula (C2) |  |  |  |  | 4.92 | 38 | 20 | 6 |
| Ventral Striatum (C2) |  |  |  |  | 4.65 | 12 | 10 | -4 |
| Pre-supplementary motor area (C3) |  |  |  |  | 4.81 | 4 | 20 | 44 |
| Pre-supplementary motor area/rostral cingulate zone (C3) |  |  |  |  | 4.19 | 8 | 30 | 30 |
| Dorsal anterior cingulate cortex (C3) |  |  |  |  | 4.11 | 8 | 36 | 20 |
| Inferior parietal lobule (C4) |  |  |  |  | 4.52 | 34 | -74 | 30 |
| Lateral intraparietal cortex (C4) |  |  |  |  | 4.46 | 40 | -40 | 44 |
| Inferior frontal cortex (C5) |  |  |  |  | 4.46 | 34 | 4 | 28 |
| Occipital cortex (C6) |  |  |  |  | 4.29 | 30 | -90 | 4 |
| Subthalamic nucleus (C7; SVC) |  |  |  |  | 4.29 | 10 | -14 | -4 |
| Ventral Striatum (C8) | 4.08 | -14 | 16 | -4 |  |  |  |  |
| Putamen (C8; SVC) | 3.89 | 30 | 18 | 0 |  |  |  |  |
| **REAL-rTMS > SHAM-rTMS** |  |  |  |  |  |  |  |  |
| *Task-related activity during “continue” trials showing a larger linear increase with cumulative gamble sum after REAL-rTMS > SHAM-rTMS covarying with change in mean stopping amount* |  |  |  |  |  |  |  |  |
| Dorsal anterior cingulate cortex | 4.13 | -6 | 8 | 32 |  |  |  |  |

Significant activation peaks (z score) and the corresponding stereotactic x, y, z coordinates in MNI space from the standard model.
Clusters are defined at $p < 0.001$ , uncorrected, at whole-brain level ( $t > 3.579$ ), cluster extent threshold 30 voxels, cluster-level $p < 0.05$ FWE-corrected. C: Cluster.

**Fig. 3.**
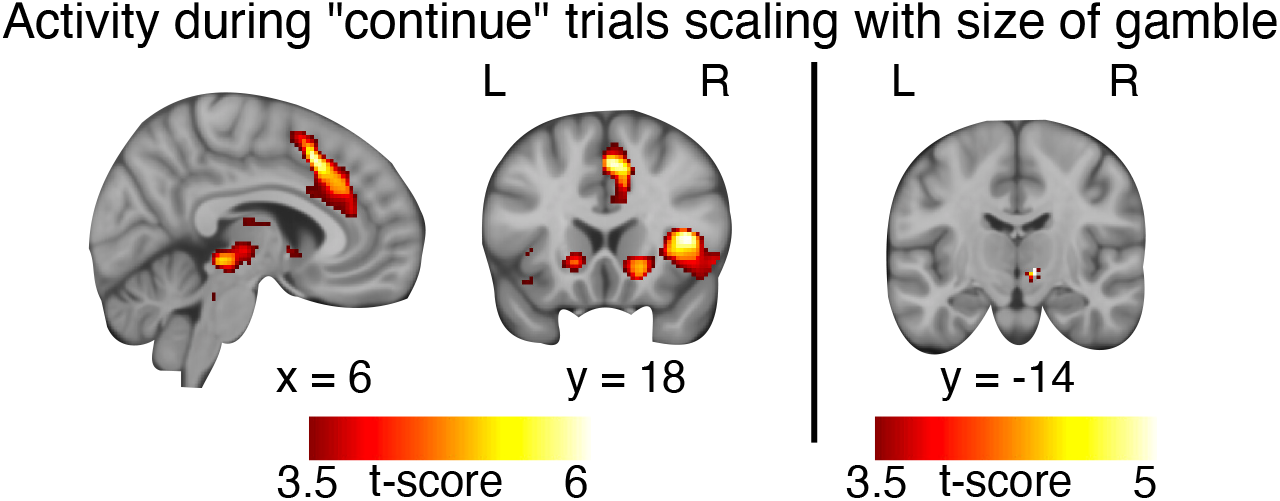
Brain regions that showed an increase in activity with the accumulated sum during “continue” trials (Table 1). *N* = 23.

The effect of real-rTMS over preSMA on choice behavior also had a functional correlate in the stimulated preSMA. The rTMS-induced change in activity in the stimulated preSMA was related to self-reported trait impulsivity (Figure 2C&D). The higher the participant’s BIS-11 score, the more did real-rTMS steepen the increase in preSMA activity with the size of the gamble (Figure 2C). The lower the participants’ BIS-11 score, the more real-rTMS flattened the rise in preSMA activity with the size of the gamble (Figure 2C). This led to a reversal of the relationship between the BIS-11 score and preSMA activation with increasing gamble size after real-rTMS relative to the sham-rTMS session (Figure 2C).

## Discussion

Here we used a perturb-and-map approach to test for a causal link between task-related activity in preSMA, impulsive choice behavior during a sequential gambling task (state impulsivity), and self-reported trait impulsivity. Using robot-assisted 1Hz rTMS, we first induced a perturbation of preSMA or performed sham stimulation. We then performed fMRI to map how real-rTMS altered regional activity in pre-SMA and impulsive choice behavior relative to sham-rTMS. We were particularly interested in assessing whether real-rTMS altered the scaling of “continue-to-gamble” activity in pre-SMA to the accumulated sum of reward during a sequential gambling round.

Our study yielded several main findings. In the unperturbed (sham-rTMS) state, self-reported trait impulsivity showed a positive linear relationship with “state” impulsivity (risky impulsivity (Robbins and Dalley, 2017)), reflected by the inclination to continue gambling for higher rewards under increasing risk. Trait impulsivity was also reflected in the task-related activation profile of preSMA in the sham-rTMS session. The lower the individual BIS-11 score, the more the preSMA increased its activity with increasing stake. Real 1Hz-rTMS of preSMA abolished these relations and the perturbation effects of real-rTMS on choice behavior and pre-SMA activity differed among individuals depending on their trait impulsivity. In the following, we first discuss the link between impulsivity and task-related preSMA activity in the absence of a preSMA perturbation. We then discuss how the perturbation of preSMA affected the relationship between impulsivity and preSMA activity in the real-rTMS session.

### Relationship between trait and state impulsivity

In the sham-rTMS session, self-reported trait impulsivity as indexed by the BIS-11 score scaled positively with risky impulsivity during the sequential gambling task (i.e., with state impulsivity probed by the task context). This is in contrast to many other studies where behavioral measures of impulsivity, including measures of stopping impulsivity, waiting impulsivity and reflection impulsivity as well as tasks applying economic notions of risk oftentimes do not reflect underlying trait impulsivity or risky behaviors in life (Cyders and Coskunpinar, 2011; Schonberg et al., 2011). While most experiments attempt to decouple different variables in order to be able to unambiguously interpret their effects on outcome measures, in this paradigm, risk and reward are coupled. This coupling is a feature of many risky decisions in real life and thus our task might elicit more naturalistic risk-taking behavior, allowing an association with trait measures of impulsivity to emerge (Schonberg et al., 2011).

### Trait impulsivity and preSMA activation with accumulated sum

Task-related fMRI showed a link between task-related activity in the preSMA and self-reported trait impulsivity. The higher the self-reported trait impulsivity, the smaller was the task-related increase in preSMA activity with increasingly risky choices. The increase in preSMA activation at higher levels of risk and rewards replicates our own previous study (Meder, Haagensen et al., 2016) and is in line with other studies with similar tasks, including the balloon analog risk-taking task (BART) (Rao et al., 2008; Helfinstein et al., 2014). Together with the preSMA, all key regions in the braking network, i.e., inferior frontal cortex, striatum and the subthalamic nucleus increase linearly with increasing stakes during continue trials. This network has been implicated in stopping behavior and conflict management (Jahfari et al., 2010; Aron et al., 2014, 2016) and the gradual build-up in activity and connectivity in this network might reflect an increasing cognitive “brake” or caution with higher sums at stake (Meder, Haagensen et al., 2016). This can be attributed to increased demands on inhibitory control, for which the preSMA has long been considered to be a key brain region (Aron et al., 2007). This finding also supports the general notion that the preSMA and dorsal medial frontal cortex are centrally involved in motivational aspects of cognitive control in general (Ridderinkhof et al., 2004; Shenhav et al., 2013; Cavanagh and Frank, 2014) and sequential value-based decision-making under risk in particular (Schonberg et al., 2012; Pearson et al., 2014). In our previous study, we found that participants with more cautious gambling behavior in the task had a stronger STN to preSMA coupling than participants who took more risky decisions (Meder, Haagensen et al., 2016). Here, we can now relate the increase in task-related preSMA activity with increasing stake to a related personality trait, impulsivity, showing that less impulsive individuals engage more effectively a neural “brake” during sequential gambling to impede risky actions in the context of increasing stakes.

### After-effects of real-rTMS of preSMA on state impulsivity and preSMA activity

Perturbing the preSMA with real-rTMS decoupled the association between trait and state impulsivity. Participants with low trait impulsivity increased their preference for risky choices after real-rTMS of preSMA, while the opposite was the case for participants with high trait impulsivity. These individuals showed a reduction in risk-taking behavior during gambling after real-rTMS.

The rTMS-induced behavioral shift was mirrored by a rTMS-induced change in task-related activity in the preSMA target. In individuals with high trait impulsivity, real-rTMS increased the otherwise low scaling of preSMA activity with the accumulated sum, whereas it decreased the otherwise higher scaling of activity in individuals with low trait impulsivity. Thus, inhibiting the preSMA had an equalizing effect on both, behavior (more risk seeking behavior in low impulsivity individuals, less risk seeking in high impulsivity individuals) and task-related preSMA activity (weaker activity increase with increasing stakes in low impulsivity individuals, stronger activity increase in high impulsivity individuals).

PreSMA is densely connected with the dorsolateral prefrontal cortex (dlPFC) and the more caudal SMA proper. One theory based primarily on fMRI data states that cognitive control of behavior is, at least in part, controlled by the prefrontal cortex organized hierarchically in a rostro-caudal order with the dlPFC on top and the cortical motor system at the bottom of the hierarchy (Badre and Nee, 2018). The preSMA is at the interface between higher-order prefrontal areas and lower-order motor areas. It has strong connections to dlPFC and the lower bank of anterior cingulate cortex (Luppino et al., 1993; Lu et al., 1994) and contains few corticospinal neurons (Luppino et al., 1994). It is, however, tightly coupled to the SMA proper and still considered a motor area (Rizzolatti and Luppino, 2001). Together with its direct connectivity with key inhibitory control regions (IFG (Luppino et al., 1993) and STN (Nambu et al., 1996)), this suggests that the preSMA might play an important role in translating strategic goals and long-term behavioral biases (traits) from prefrontal areas into specific adaptive actions implemented by the motor system. We speculate that by perturbing the activity in preSMA the rostro-caudal chain of information might be impaired, leading to a decoupling of long-term, abstract goals, represented in the more rostro-lateral prefrontal cortex and the motor implementation in the cortical motor system. This might lead to the decoupling of trait impulsivity and state impulsivity, i.e., the task-related choice behavior.

This baseline dependency is an effect oftentimes observed in the context of neuromodulatory neurotransmitters such as dopamine or noradrenaline, where too low and too high levels of the neurotransmitter lead to poorer task performance (Robbins and Arnsten, 2009; Dayan, 2012; Meder et al., 2018). It is often referred to as the Yerkes-Dodson function or an inverted U-shape (Robbins, 2000). Perturbing the preSMA might release the region from the effects of prefrontally mediated “too low” and “too high” levels of trait impulsivity which otherwise modulate the area’s response to risky decisions and the resulting choice behavior. Personality traits might be associated with different baseline levels of neural activity in the preSMA which may result in bidirectional effects of rTMS depending on the level of activity such that high activity level leads to suppression and low activity level leads to facilitation of neural activity reflecting the principle of homeostatic plasticity (Siebner, 2004). This moves activity and behavior towards the highest point of the inverted U-shape. Note, however, that real-rTMS did not actually lead to more optimal stopping behavior in the sense of gain-maximization, i.e. the mean stop amount was not closer to the gain-maximizing 200 DKK. Optimal behavior should here thus be understood in terms of utility where risk- and loss-aversion are commonly observed in gambling decisions (Kahneman and Tversky, 1979).

To our knowledge this is the first study to show that preSMA is centrally involved in translating trait impulsivity into actual choice behavior. Furthermore, we provide evidence that the functional significance of preSMA activity for choice behavior under risk depends on the trait impulsivity of the individual. In future studies, perturbation of preSMA in populations with excessive levels of impulsivity such as pathological gambling or substance abuse disorder could help gain mechanistic insight and potential treatment options for these complex and disabling disorders.

## Acknowledgments

This work was supported by the Danish Parkinson Foundation (Parkinsonforeningen), The Capital Region of Denmark (Region Hovedstaden), Amager and Hvidovre Hospitals, Bispebjerg and Frederiksberg Hospitals and The Danish Movement Disorder Society. D.M. was supported by the Novo Nordisk Foundation (Grant No. NNF16OC0023090). H.R.S. was supported by a Collaborative Alliance grant from Lundbeck Foundation (R336-2020-1035). H.R.S. holds a 5-year professorship in precision medicine at the Faculty of Health Sciences and Medicine, University of Copenhagen, which is sponsored by the Lundbeck Foundation (Grant No. R186-2015-2138). None of the funding sources were involved in the undertaking of the study.

